# Neuronal dynamics of signal selective motor plan cancellation in the macaque dorsal premotor cortex

**DOI:** 10.1101/2020.01.08.897397

**Authors:** Giarrocco Franco, Bardella Giampiero, Giamundo Margherita, Fabbrini Francesco, Brunamonti Emiliano, Pani Pierpaolo, Ferraina Stefano

## Abstract

Primates adopt various strategies to interact with the environment. Yet, no study has examined the effects of behavioral strategies with regard to how movement inhibition is implemented at the neuronal level. We modified a classical approach to study movement control (stop-task) by adding an extra signal – termed the Ignore signal – which influenced movement inhibition only under a specific strategy. We simultaneously recorded multisite neuronal activity from the dorsal premotor (PMd) cortex of macaque monkeys during a task and applied a state-space approach. As a result, we found that movement generation is characterized by neuronal dynamics that evolve between subspaces. When the movement is halted, this evolution is arrested and inverted. Conversely, when the Ignore signal is presented, inversion of the evolution is observed briefly and only when a specific behavioral strategy is adopted. Moreover, neuronal signatures during the inhibitory process were predictive of how PMd processes inhibitory signals, allowing the classification of the resulting behavioral strategy. Our data corroborate the PMd as a critical node in movement inhibition.

## 1 Introduction

Primates have the crucial ability to suppress actions rapidly, a capacity that can be strategically regulated. For example, they can decide to procrastinate an ongoing action if something occurs suddenly in the environment: this momentary pause could allow them to better evaluate the consequence of the action and decide their next move. Alternatively, they can choose to ignore the new signal and continue pursuing the initial goal.

Neuroscientific studies have typically examined simple forms of movement inhibition in experimental settings, using the stop (or countermanding) task (Logan and Cowan, 1984; Hanes et al., 1998). In this task, the primary instruction is to respond as quickly as possible to a Go signal (no-signal trials); in a minority of trials (stop-signal trials) a Stop signal is presented after the Go signal, and subjects are required to refrain from moving. The ability to countermand the response is evaluated, based on the estimate of the stop signal reaction time (SSRT). The SSRT can be broadly considered to be the response time of movement inhibition (Logan and Cowan, 1984) and as such has been used to compare the efficiency of inhibitory control in various populations of patients; during the many stages of brain maturation, from childhood to senescence; and across animal species (Pani et al., 2013; Brunamonti et al., 2011; Brunamonti et al., 2012; Lipszyc and Schachar, 2010; Lijffijt et al., 2005; Hippolyte et al., 2009; Williams et al., 1999; Hanes et al., 1998; Paré and Hanes, 2003; Pani et al., 2014).

However, because inhibition can intervene to regulate behavior in many ways, in recent years, more complex tasks have been used to determine the behavioral consequences of and the neuronal functional architecture that underlies complex inhibitory control (Bissett and Logan, 2014; Sebastian et al., 2017; Xu et al., 2017; Xu et al., 2018; Aron, 2010; Sharp et al., 2010; Boehler et al., 2011; Cai et al., 2011; Chikazoe et al., 2009; Majid et al., 2012; Sebastian et al., 2016). Among the tasks that have proposed, the **selective stop task** (Bissett and Logan, 2014) entails the presentation of a Stop signal or an Ignore signal after the Go signal in a subset of trials, of which only the Stop signal requires inhibition of the movement.

An important feature is that subjects can use several strategies to solve the task when the Ignore signal is presented; each strategy requires a different method to implement inhibitory control. Subjects can choose to first discriminate between the Stop/Ignore signals and then decide to hold or move; alternatively, they can momentarily inhibit the response, irrespective of the signal, which appears after discrimination of the Stop/Ignore signal, to generate or cancel the movement. In the literature, the definition of the adopted strategy is based strictly on the analysis of the behavioral data; it is unknown whether the Ignore signal, when behaviorally relevant, temporally activates the inhibitory neuronal process.

The level of response control that is required in stop tasks is supported by a network of cortical and subcortical brain regions; nonetheless, the specific function of each node continues to be investigated (Cai et al., 2011; Aron et al., 2007; Wessel and Aron, 2016; Aron et al., 2006; Aron et al., 2003; Wagner et al., 2018; Duque et al., 2012; Duque et al., 2017; Parmigiani and Cattaneo, 2018; Hanes et al., 1998; Paret’ and Hanes, 2003; Schmidt et al., 2013; Mallet et al., 2016). Of the various cortical regions, the dorsal premotor cortex (PMd) appears to be crucial in establishing the transition from decision processes to the action of reaching movements: neuronal activity in this area continuously reflects the accumulation and change in information that is pertinent to the momentary decision state regarding forthcoming movements (Thura and Cisek, 2014; Kaufman et al., 2014), and predicts when and whether an arm movement will be generated or inhibited (Kaufman et al., 2016; Mirabella et al., 2011; Pani et al., 2018; Pani et al., 2014). Evidence suggests that neuronal modulation in the PMd contributes to the inhibition of movements disparately under different strategies and, consequently, that the PMd is an ideal region to examine the various aspects of the underlying neuronal dynamics with the high temporal definition of neurophysiological approach.

In this study, we determined whether and how the Ignore signal influences the neuronal activities that are related to preparation of the motor plan in the PMd. By analyzing the multisite dynamics of neuronal activity in the PMd, we obtained evidence of a strategy-dependent inhibitory neuronal process, triggered by the Ignore signal. These novel data demonstrate the strong correlation between modulation of the neuronal activity in the PMd and a signal-specific decision for inhibition, further demonstrating that the PMd is a key structure in movement control.

## 2 Methods

### 2.1 Subjects

Two male rhesus macaque monkeys (Macaca mulatta; designated Monkeys 1 and 2), weighing 9 and 13 kg, were studied. The monkeys were pair-housed with cage enrichment. They were fed daily with standard primate chow, supplemented with nuts and fresh fruits. The monkeys received part of their daily water supply during the experiments in the form of fruit juice. All experimental procedures, animal care, housing, and surgical procedures conformed to European (Directive 2010/63/UE) and Italian (D.L. 26/2014) laws on the use of nonhuman primates in scientific research and were approved by the Italian Ministry of Health.

### 2.2 Animal preparation

At the end of training period a Utah array (96 channels, Blackrock Microsystems, USA) was implanted in the PMd of each monkey, using the the arcuate sulcus (AS) and pre-central dimple (pCD) as anatomical landmarks after opening of the dura (Fig. 1 in Supplementary materials). The site of the implant was contralateral to the arm that was used during the experiment. All surgeries were performed under sterile conditions and veterinary supervision. Antibiotics and analgesics were administered postoperatively. Anesthesia was induced with ketamine (Imalgene, 10 mg kg^−1^ i.m.) and medetomidine hydrochloride (Domitor, 0.04 mg kg^−1^ i.m. or s.c.) and maintained with inhalant isoflurane (0.5% to 4%) in oxygen. Antibiotics were administered prophylactically during the surgery and postoperatively for at least 1 week. Postoperative analgesics were given at least twice daily. Recordings of neuroanl activity started after a minimum of 10 weeks after recovery from the surgery. A head-holding device was implanted in Monkey 2 before the training started, whereas it was implanted simultaneously with the array in Monkey 1.

**Figure 1:**
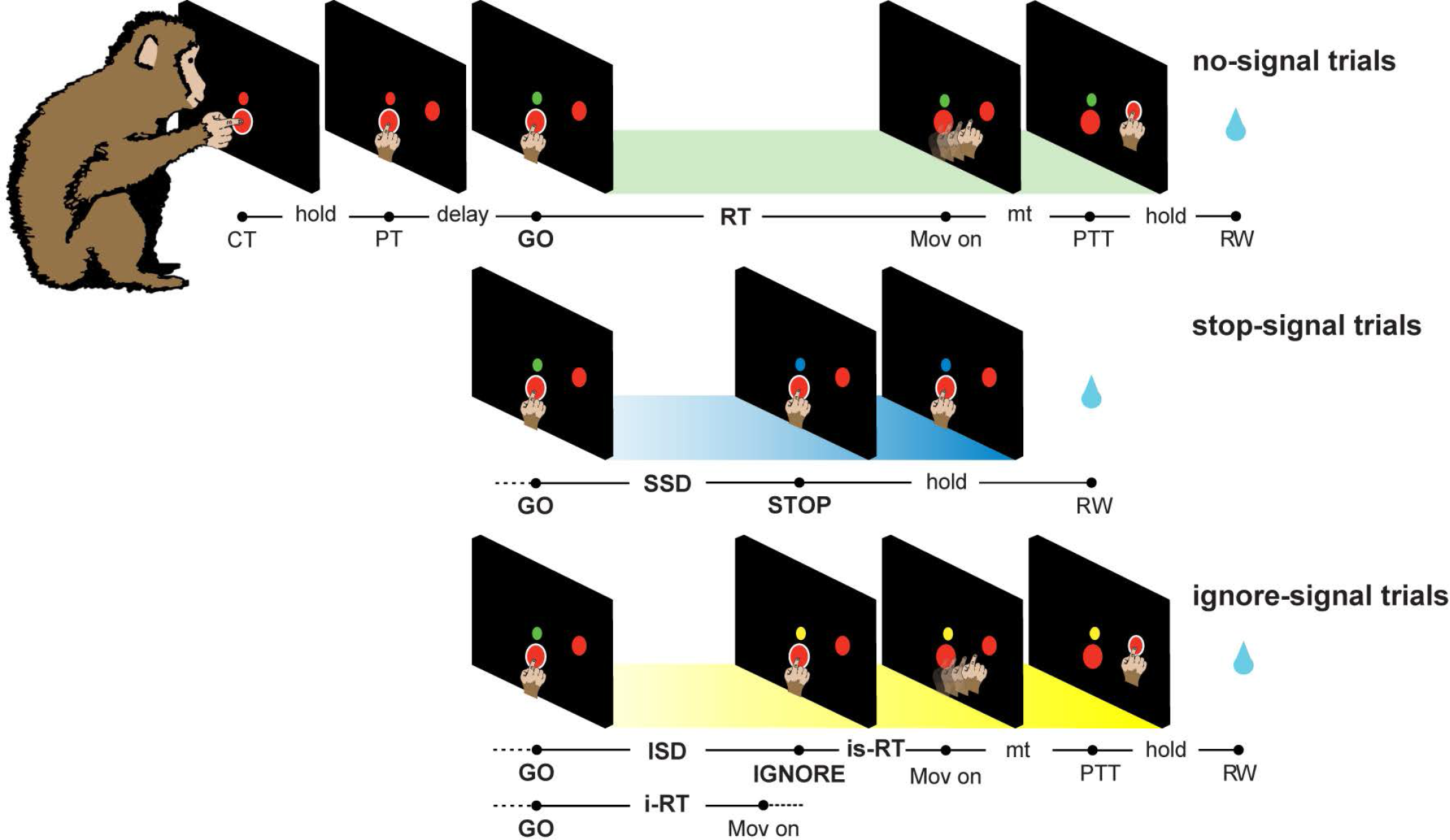
Signal selective stop task. Three types of trials were intermingled in each block. Movement cancellation was required in stop-signal trials only. CT, central target; PT, peripheral target; GO, Go signal; Mov on, movement onset; RT, reaction time; is-RT, Ignore-signal reaction time; i-RT, ignore reaction time; mt, movement time; PTT, peripheral target touch; RW, reward. SSD, stop signal delay; ISD, ignore signal delay; hold, central, or peripheral holding time.

### 2.3 Apparatus and task

Experiments were performed in a darkened, acoustically insulated room. Monkeys were seated in front of a black isoluminant background (<0.1 cd/m^2^) of a 17-inch touchscreen monitor (LCD, 800 × 600 resolution). A noncommercial software package, CORTEX (www.nimh.nih.gov), was used to control the presentation of stimuli and behavioral responses. Figure 1 shows the schema of the task, comprising 3 types of trials, randomly intermixed: no-signal trials (60%), stop-signal trials (20%), and ignore-signal trials (20%). Each trial started with the appearance of a central target (CT) (red circle, diameter 1.9 cm) and a Cue signal (red circle, diameter 0.7 cm), smaller and appearing slightly above (3 cm) the CT. The monkeys had to touch the CT and keep their finger on it for various holding times (500-800 ms, 50-ms step). Subsequently, a peripheral target (PT) (red circle, diameter 1.9 cm) appeared randomly at 1 of 2 possible locations (i.e., 7 cm to the right or left of the vertical midline of the screen; right only in 4/5 sessions Monkey 2). In all trials, after a fore-period delay (variable duration, depending on the session), a Go signal appeared, consisting of a green circle, replacing the Cue (stimulus circle 0.7 cm RGB: [0 250 0]; 85 cd/m^2^). The Go signal instructed the subjects to react and reach for the PT as quickly as possible. In no-signal trials, the task continued, and after the movement onset, the animal was required to maintain its touch at the new position for a variable time (600-800 ms, 50-ms step) until the end of the trial. The RT (reaction time) was defined as the time between the presentation of the Go signal and the onset of the hand movement. An upper temporal limit (upper RT) was set for no-signal trials to 1200 ms. This value was set gradually during the training to instruct the monkey to respond quickly, avoiding waiting for the (potential) Stop signal. In the stop-signal and ignore-signal trials, the sequence of events was the same until the presentation of the Go signal. In stop-signal trials, the Stop signal (blue circle, 0.7 cm RGB: [0 0 188], 7 cd/m2) replaced the Go signal at the end of a variable and unpredictable interval (stop-signal delay, or SSD). In these trials, a hand that was kept on the CT until the end of the trial (800-1000 ms, 50-ms step) corresponded to a correct response (signal-inhibit trials). Conversely, the simple detachment of the hand after the presentation of the Go signal was defined as an incorrect response (signal-respond trials).

Similarly, in ignore-signal trials, the Ignore signal (yellow circle, 0.7 cm RGB: [255 255 0], 65 cd/m^2^) replaced the Go signal after a variable interval (ignore-signal delay or ISD). In these trials, the monkeys were instructed to respond as they did during no-signal trials, discounting the Ignore signal. We defined ignore-RT (i-RT) as the time between the presentation of the Go signal and the onset of the hand movement, regardless of the appearance of the Ignore signal. If the onset of the hand movement occurred during the ISD (i.e., before the appearance of the Ignore signal), the trial was still considered to be a correct trial. Further, we defined ignore-signal RT (is-RT) as the time between the presentation of the Ignore signal and the onset of the hand movement. No upper RT was set in ignore-signal trials. At the end of correct trials, the monkeys experienced a brief sound that was accompanied by the delivery of juice as a reward. In signal-respond trials (errors), neither sound nor reward was delivered, and the screen turned blank. The intertrial interval was set to 1000 ms. Various fore-period delays (ranging from zero ms, no-delay, to 1150 ms) were used in different sessions (for further details, see Table 1, supplementary materials). Our purpose was to affect the level of movement preparation and the strategy that was used consequently. In humans, subjects adopt different strategies in the Selective stop task by manipulating other factors that influence the movement preparation, such as the relative proportion of stop versus ignore trials (Bisset and Logan 2014). During stop-signal trials, a staircase procedure was adopted to determine the duration of the SSDs for each trial as follows: if the monkey succeeded in withholding the response (signal-inhibit trials), the SSD increased by 1 step (100 ms for both monkeys) in the subsequent stop-signal trial; conversely, if the subject failed (signal-respond trials), the SSD decreased by 1 step. The SSDs ranged from 120 ms to 1020 ms, depending on the performance. For each ignore-signal trial, the ISD was set to the SSD in the previous stop-signal trial. The same procedure has been adopted in previous studies on human subjects (Bissett and Logan, 2014).

### 2.4 Behavioural analysis

We analyzed the monkeys behavioral performance in the framework of the race model (Logan and Cowan, 1984). The race model assumes that during stop-signal trials, 2 stochastic processes race toward a threshold — the GO and STOP processes — triggered by the appearance of the Go and Stop signals, respectively. The result of this race — movement generation (signal-respond trials) or movement inhibition (signal-inhibit trials) — depends on which of these processes reaches its threshold first. In signal-inhibit trials, the STOP process wins over the GO process, and vice versa in signal-respond trials. The main assumption of the race model is that the GO and STOP processes are independent of each other (independence assumption). In particular, the model assumes 2 types of independence: first, that in a given trial, the latency of the GO process does not depend on the latency of the STOP process (stochastic independence), and second, that the GO process in the stop-signal trials must be the same as in the no-signal trials, because the GO process must be unaffected by the presence of the Stop signal (context independence). Practically, to validate the independence assumption, signal-respond trial RTs must be shorter than the RTs for no-signal trials (Logan and Cowan, 1984). Only if the independence assumption is validated can the race model be used to estimate the SSRT in stop-signal trials by setting 3 variables: the no-signal trials RTs, the probability of responding (or error) to the Stop signal, and the duration of the SSD. We used the integration method to obtain the SSRT. This method assumes that the latency of the STOP process corresponds to the nth no-signal RT, where n results from the mathematical product of the distribution of ordered no-signal RTs and the overall probability of responding when using the tracking procedure. The SSRT can then be calculated by subtracting the mean SSD from the nth no-signal RT (Band et al. 2003).

In the selective stop task, we assigned a behavioral strategy to each session as previously proposed (Bissett and Logan, 2014; Sebastian et al., 2017) by comparing no-signal RTs, signal-respond RTs, and ignore-signal RTs. The analysis of all of the sessions confirmed the independence assumption – ie, signal-respond RTs were faster than no-signal RTs (see Results). A **stop-then-discriminate (STD)** strategy was assigned if the ignore-signal RTs were slower than no-signal RTs. Conversely, a **discriminate-then-stop (DTS)** strategy was assigned if the ignore-signal RTs were not slower than no-signal RTs. The rationale for these strategies is as follows: in the STD strategy, the subjects suppress their responses on the appearance of a signal that follows the Go signal and then restart the movement process after detecting it as an Ignore signal; in the DTS, the subjects first discriminate between signals (Ignore vs Stop), suppressing the response only on detection of a Stop signal. To provide statistical support for the assignment of categories, we performed t-test between the RTs of various types of trials and then applied Jeff Rouders Bayes factor calculator (http://pcl.missouri.edu/bayesfactor)-ie, we converted t-value and sample size to a Bayes factor, thereby obtaining the odds in favor of the null or alternative hypothesis. We analyzed the distribution of ignore-signal RTs by Hartigans Dip Test of Unimodality. Whenever a non-unimodal distribution was observed (in all cases, bi-modal distribution), we evaluated the time when the separation between curves occurred by k-means clustering method for the ordered ignore-signal RTs.

### 2.5 Neural recordings and analysis

Unfiltered raw activity was recorded from 96 channels using a Utah array (Blackrock Microsystems, USA) and a TDT System 3 (Tucker Davies Technologies, sampling rate 24.4 kHz). From the raw signal that was recorded from each channel (site), we extracted the spectral estimate of the multiunit activity (MUA) offline as a good approximation of the average firing rate, as described in (Mattia et al., 2013). The MUA was smoothed using a moving average sliding window (±20-ms sliding window, 5-ms step).

From each recording session, we selected the sites with significant difference (t-test, p< 0.05) between baseline (from 200 to 400 ms following the touch of the CT) and the activity during at least 1 of 3 relevant trial events (from 250 to 50 ms before the movement onset in no-signal trials; the 200 ms following the Stop signal onset in stop-signal trials; and the 200 ms following the Go signal in no-signal and stop-signal trials). We considered the activity that was recorded from these sites to be **task-related**. We then compared the activity between signal-inhibit trials and latency-matched no-signal trials. The latter comprised no-signal trials in which RTs were longer than the sum of the average SSD and the SSRT that was computed in the same session (ie, in agreement with the race model assumptions) and had a similar level of movement preparation as in signal-inhibit trials. In signal-inhibit trials, the activity was aligned to the Stop presentation, and in latency-matched no-signal trials, it was aligned to the hypothetical presentation of the Stop signal (ie, Go signal presentation + SSD). To determine whether a difference in MUA occurred between these 2 trial types, we applied the shuffle test for each interval, from 50 ms before to 250 ms after the onset of the Stop signal — a method previosuly used in other studies on movement inhibition (Mallet et al., 2016; Schmidt et al., 2013). First, for each interval, we compared the mean of the 2 trial types. Then, we shuffled the 2 trial types 10,000 times for each interval, and for each shuffle, we compared the means of the 2 resulting trial distributions. We acquired a P value by counting the number of shuffles in which the difference between the obtained means was larger (or smaller) than that between the 2 observed means. We used a P value of 0.05 to define whether the difference between signal-inhibit trials and latency-matched no-signal trials was significant. Thus, for each interval, we considered only sites for which were more than 9500/10000 different means. Then, we performed the binomial test (p=0.05) to set a threshold and determine whether the measured fraction of sites was statistically significant for each interval.

We replicated the shuffle test for each pair of trial types to quantify the fraction of sites that distinguished between the 3 types of trials according to the behavioral strategy. When the ignore-signal trials were included in the shuffle test, the activity was aligned to the presentation of the Ignore signal.

At the end of this analysis, we selected the sites with activity that **participated to inhibitory control** — ie, sites with activity that exceeded the baseline threshold for 50 ms during the SSRT when contrasting latency-matched no-signal with signal-inhibit trials — and proceeded to the next analyses.

### 2.6 Principal components analysis (PCA)

To describe the multisite neuronal dynamics that were related to various behavioral strategies, we transformed the neuronal data by principal component analysis (PCA). We created a matrix N (recording sites) × P (MUA time value) by concatenating the average MUA across no-signal, ignore-signal, and signal-inhibit trials for each site. The MUA was aligned to the onset of the Stop signal (signal-inhibit trials), Ignore signal (ignore-signal trials), and a hypothetical Stop signal (latency-matched no-signal trials). The MUA spanned the interval from the average presentation of the Go signal to 800 ms following the time of alignment. We represented the evolution of neuronal dynamics as neuronal trajectories in a 3D state space, in which the 3 dimensions were the first 3 principal components that explained most of the variance (at least 90%). We reported the analysis separately for each recording session, with 1 exception: we combined the activity of 2 sessions from Monkey 2 when the DTS strategy was adopted, to increase the sample size and the reliability of the data. In these sessions, to avoid oversampling from the same population of neurons, we selected only activity derived from different sites or, when from the same site, activity with different patterns in relation to movement generation and inhibition (see Supplementary Fig. 2). To quantify the divergence between neuronal trajectories across trials that were related to movement inhibition, we measured the temporal evolution of the first derivative of the projection of the neuronal state onto the single dimensions. For each dimension, we first defined the **Signal Neuronal Time** (SNT) as the time in which the evolution of the projection changed direction after the Stop signal (signal-inhibit trials) or the Ignore signal (ignore-signal trials) — ie, the time in which the derivative was equal to 0. Then, we calculated the weighted average of the resulting SNTs, based on the weight of the explained variance of the single dimensions.

### 2.7 Relationship between neuronal activity and behavioral strategy

To tentatively predict the behavioral strategy from neuronal activity modulations, we characterized how the MUA differed when the movement would have been generated under different conditions (no-signal and ignore-signal trials) with respect to when it would have been inhibited (signal-inhibit trials). To this end, we implemented an algorithm, based on formula (1), designed to perform the following comparisons: signal-inhibit trials versus latency-matched no-signal trials and signal-inhibit trials versus ignore-signal trials (ignore-signal trials with RTs>ISD+SSRT). The inputs into the algorithm were the MUA aligned to the Stop signal (signal-inhibit trials), Ignore signal (latency-matched ignore-signal trials), and average SSD value (latency-matched no-signal trials). The MUA interval spanned from −50 ms to +225 ms relative to the time of alignment. This interval always occurred before the behavioral strategy can be defined — ie, before RTs can be detected. Moreover, the same types of trials were selected from each session with the same criteria, and the selection process was not affected by the behavioral strategy.

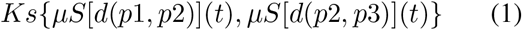

For each of the 2 comparisons, the algorithm performed the following steps. From each trial type, it randomly selected trials 1000 times. Because each category of trial had a different number of trials, the algorithm was set to select 75% of the total number of trials in the smallest category. Then, for each site and interval, it computed the squared Euclidean distance (d) of the activities between different categories of trials (p1, latency-matched no-signal trials; p2, signal-inhibit trials; p3, ignore-signal trials), obtaining an ensemble of 1000 squared Euclidean distances for each comparison.

This first step provides an evolution of the squared Euclidean distances over time — ie, a measure of how the activities between the 2 selected categories of trial differ over time. This step was repeated for all sites, and a mean Euclidean distance was obtained by averaging the values across all recording sites. The output was an n (1000 averaged squared Euclidean distance across sites) × m (squared Euclidean distance time values) matrix, in which the slope of each of the 1000 distances, fitted every 50 ms with a step of 5 ms, was calculated. This step provides an evolution of the distribution of slopes, used to quantify how the difference between activities changed over time (t). Finally, each resulting slope distribution was averaged, obtaining a single averaged slope distribution (μS) from -25 ms to +200 ms relative to the signal presentation, for each comparison. In the last step, Kolmogorov-Smirnov test (KS) was performed between the 2 resulting slope distributions. To confirm the presence of 2 separate sets of p values, we also performed cluster analysis, based on the distances between the observed p values.

## 3 Results

### 3.1 Behavioral results

We derived behavioral data from 15 sessions (see Supplementary Materials: Table 1; Monkey 1, 10 sessions; Monkey 2, 5 sessions) with a sufficient number of trials for analysis. Monkeys adopted the DTS or STD strategy on different days.

Consistent with the independence assumption of the race model, in both strategies, the SSRT divides the distribution of RTs in no-signal trials (Fig. 2, A shows 2 sample sessions; 1 for each strategy for the same animal) for fast (left portion) and slow (right portion) responses. As expected, signal-respond RTs overlapped primarily with fast responses in both strategies and are thus shorter than no-signal RTs (Fig, 2, B; Table 1 in Supplementary Materials). Conversely, the distribution of responses in the ignore-signal trials showed strategy-specific differences. In the same sample session, the histograms in the lower area (Fig. 2, D) show that in the STD strategy, a bimodal distribution emerged in the is-RTs [p(max) <.001]. This pattern suggests that in certain trials (red bars), the Ignore signal activated behaviorally relevant inhibitory motor control, as confirmed by the longer duration of i-RTs compared with no-signal RTs (Fig. 2, C). Similar results were observed in all STD sessions (Monkey 1, 6 sessions: i-RTs: mean 941 ms, SD 170; no-signal RTs: mean 906.7 ms, SD 151.4, p<.001, rank-sum test; Monkey 2, 1 session; see Supplementary Materials: Table 1 and Supplementary Fig. 3). The slow mode of is-RTs always resulted in their occurring after the SSRT (Fig.2, D), strengthening the hypothesis that when the monkeys adopt the STD strategy, they first inhibit and then restart the response after discrimination of the Ignore signal. When the monkeys adopted the DTS strategy during the ignore-signal trials, we did not find any behavioral signs of inhibitory control that was driven by the Ignore signal (Fig. 2 for a sample session; see Fig. 3 in Supplementary Materials). None of the is-RT distributions showed bi-modality [p(min) > 0.7], and further, we did not observe a longer duration of i-RTs compared with no-signal trial RTs (Monkey 1, 4 sessions: i-RTs mean 676 ms, SD 197.2; no-signal RTs mean 683.1 ms, SD 177.9, p=0.35; Monkey 2, 4 sessions: i-RTs mean 580 ms, SD 235.2; no-signal RTs mean 575 ms, SD 218.1, p=0.58, rank-sum test).

**Figure 2:**
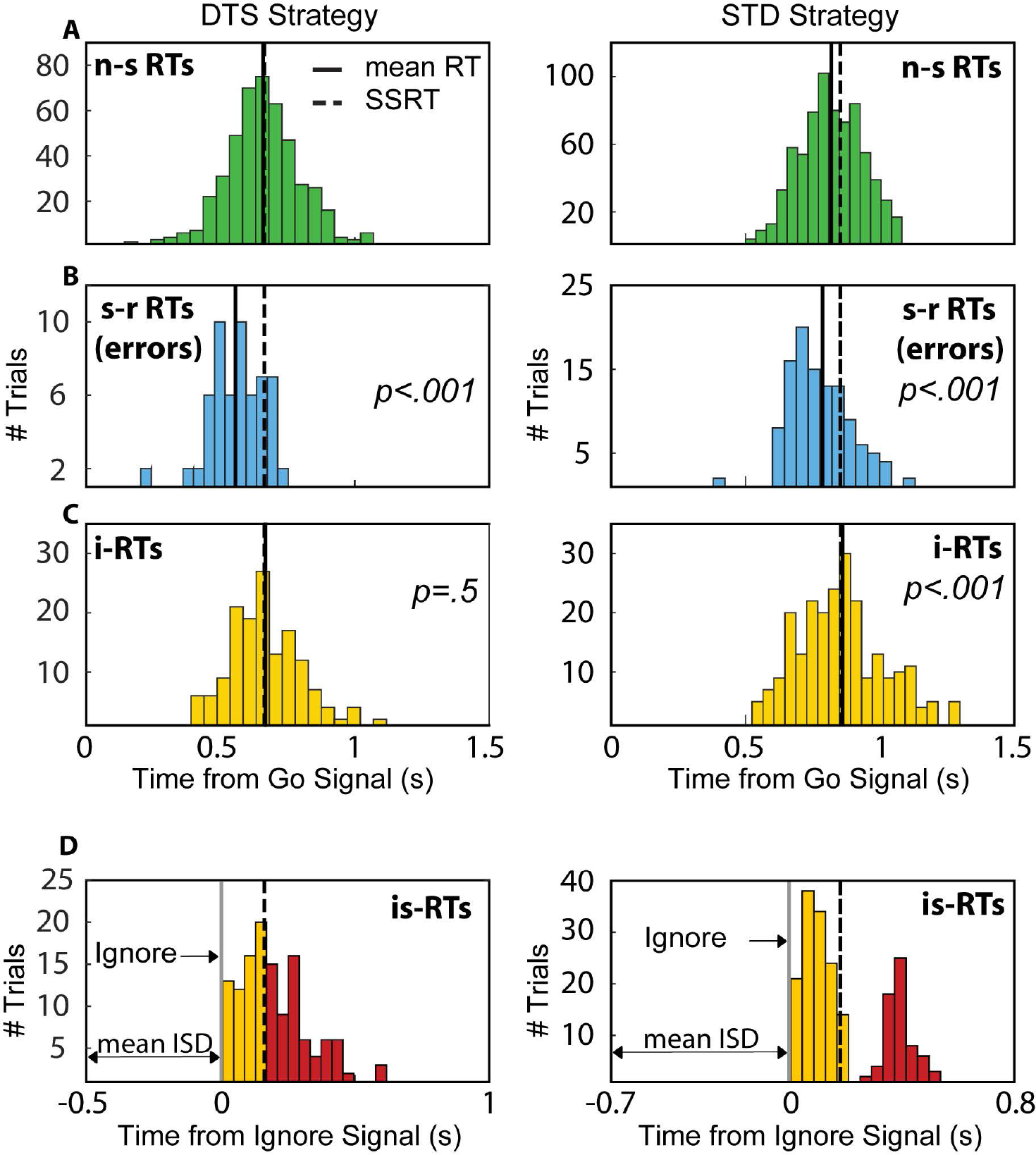
Reaction time distributions for both strategies. Each column shows data for sessions classified by one of the two strategies (see Methods). A, n-s, no-signal trial RTs. B, s-r, signal-respond trial RTs. C, ignore-signal trial RTs. In B and C, the P values are for the comparison (rank-sum test) with data in A. Continuous line: average RT; interrupted line: SSRT. D, same data in C aligned to the Ignore Signal (Ignore) and sorted by duration (in DTS, red bars are is-RTs > SSRT; in STD, red bars are those in the slow mode of the bi-modal distribution).

**Figure 3:**
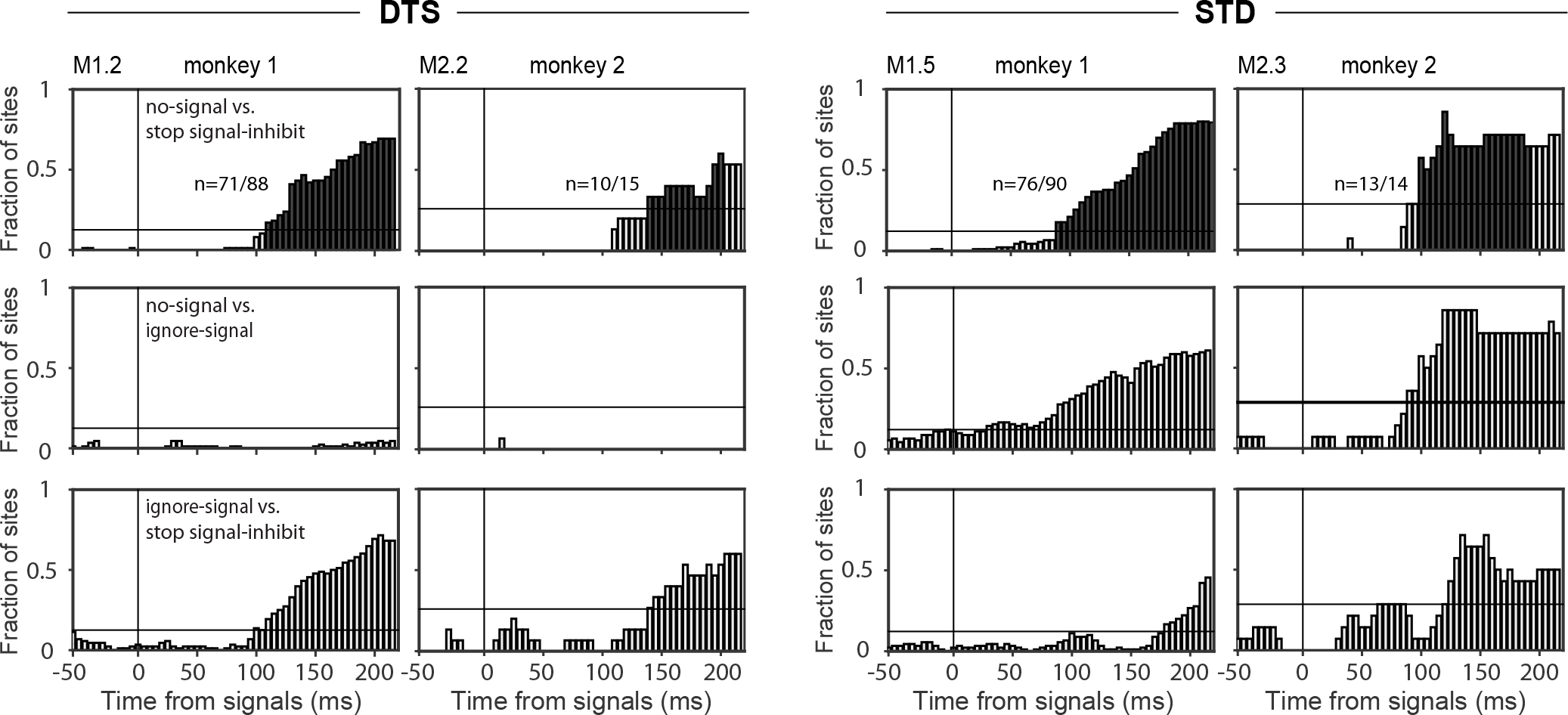
Neuronal characterization of behavioral strategies at the population level. Fraction of sites with significantly different MUAs for the trial types under comparison. Data are presented separately for each monkey and for the two strategies (DTS and STD). We compared latency-matched no-signal, signal-inhibit, and ignore-signal trials (see colored bars at the bottom for the specific comparison). The alignment time is the Stop signal for signal-inhibit trials, the Ignore signal for ignore-signal trials, and the hypothetical presentation of the Stop signal for latency-matched no-signal trials. The analysis was performed across all sites showing task-related activity (mumbers are in panels at the top of each column). The horizontal line on each plot corresponds to the threshold obtained after shuffle test (p<0.05) between each pair of trials and the binomial test (p=0.05) (see Methods). Dark bars in the top plots indicate the fraction of sites that significantly exceeded the threshold during the SSRT (see Methods).

Following the observation that the Ignore signal influences behavior only when the STD strategy is adopted, we examined whether the length of the inhibitory process in stop-signal trials is affected by the behavioral strategy (see Table 1 in Supplementary Materials). We did not observed any differences between SSRTs as a function of behavioral strategy (Monkey 1, STD strategy: mean SSRT=214.67 ms, CI=[201.92, 227.41], DTS strategy: mean SSRT=205.25 ms, CI=[176.92, 233.58], rank-sum test p=0.53; Monkey 2, STD strategy: SSRT=203 ms, DTS strategy: mean SSRT=204.50 ms, CI=[176.14, 232.86]); the same results were obtained after combining the data from both monkeys (STD strategy: mean SSRT=213 ms, CI=[201.97, 224.03]; DTS strategy: mean SSRT=204.88 ms, CI=[191.08, 218.67], rank-sum test p=0.30). The behavioral definition of strategies is based on the decision matrix per Logan and Bisset (2014); when computing other metrics (see Supplementary Materials Fig. 4), the distinction remains, although a smoother transition between the 2 strategies can be observed.

**Figure 4:**
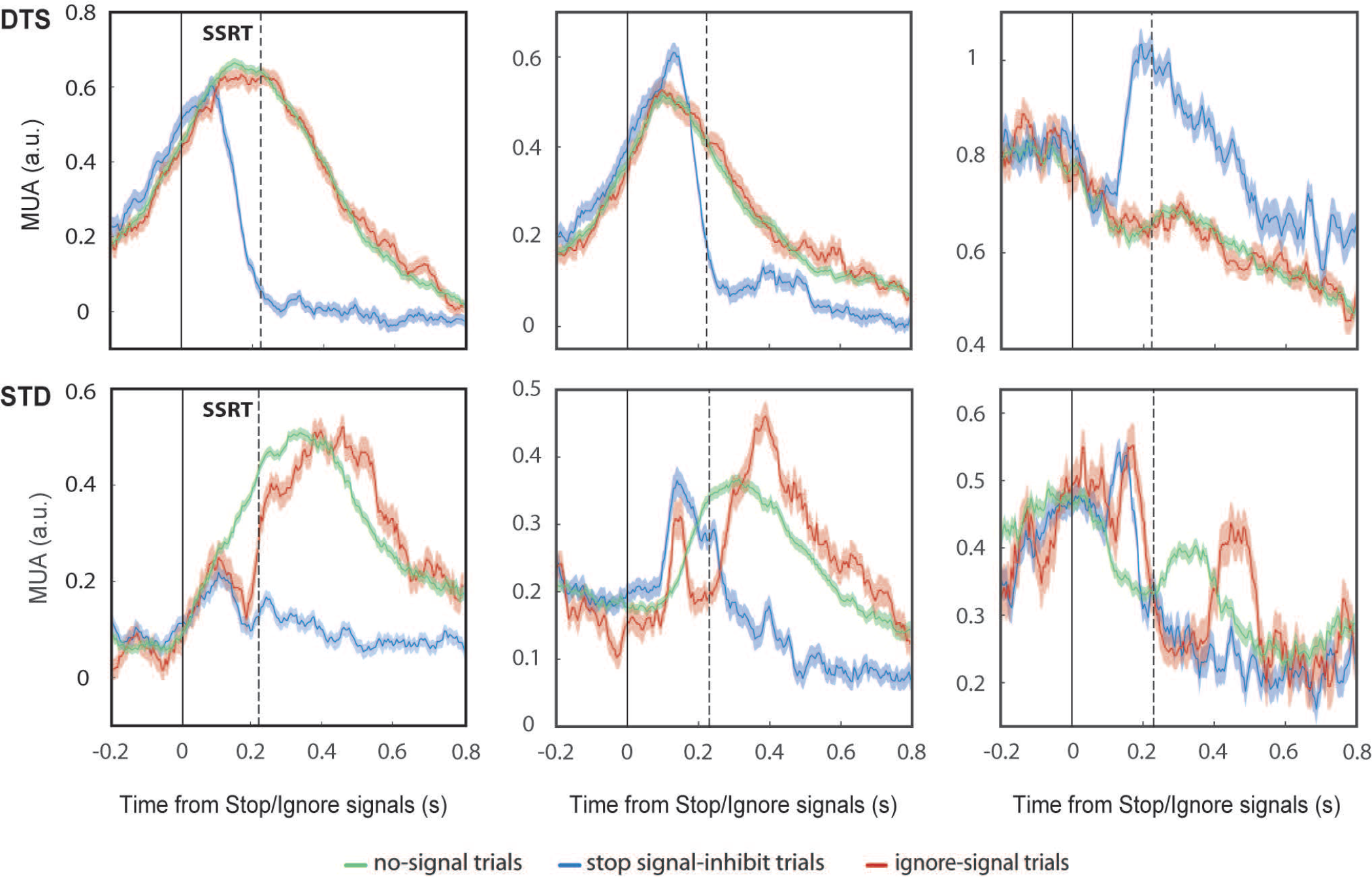
Examples of neuronal patterns for the 2 behavioral strategies. Average MUA (±SE) from three sample sites are shown for each trial type and strategy (DTS and STD). sites are examples of those extracted by the shuffling analysis depicted in Figure 3. SSRT: stop signal reaction time.

We were unable to demonstrate a clear link between fore-period duration and strategy. For example, Monkey 2 used an STD strategy with long fore-period delays, and Monkey 1 used a DTS strategy with zero delay (see Table 1, supplementary materials).

In conclusion, the behavioral evidence suggests that the Ignore signal temporarily inhibits behavior only when the STD strategy was adopted. Concurrently, the inhibitory process that is driven by the Stop signal does not appear to be influenced by the strategy.

### 3.2 Neuronal results

We were able to record reliable neuronal signals in 8 sessions. In the other sessions (n=7), technical problems prevented us from adequately collecting raw signals to analyze. To test whether the Ignore signal differently influences the neuronal inhibitory process according to the behavioral strategy, we first identified the sites that experienced neuronal modulation that could be involved in movement inhibition. Figure 3 (top row) shows the fraction of sites that discriminated between trials when the movement was generated (latency-matched no-signal trials) versus inhibited (signal-inhibit trials) in 4 representative sessions - 1 for each monkey and strategy (see Supplementary Fig. 5, for other sessions). Across sessions and monkeys, approximately 78% of task related sites (mean ±SE: 80 ±4% Monkey 1 and 73 ±10% Monkey 2; p<0.05, shuffle test) showed a significant (p<0.05, binomial test) difference between no-signal and signal-inhibit trials during the SSRT (Figure 3, black bars; see Methods for further details). Further, we evaluated the fraction of task related sites that modulated differently between trial types, based on the behavioral strategy. We replicated the shuffle test, comparing the activity in ignore-signal trials with that in latency-matched no-signal trials and signal-inhibit trials respectively (Figure 3, middle and bottom rows). Significantly different modulation during the DTS strategy typically occurred when ignore-signal trials were compared with signal-inhibit trials. A small fraction of sites was differently modulated in no-signal and ignore-signal trials, supporting the presence of similar neuronal modulation when the movement was generated under these two conditions. The test differed when the STD strategy was used. Similar patterns of discrimination occurred in comparing latency-matched no-signal trials with signal-inhibit and ignore-signal trials, demonstrating that in the STD strategy, the MUA that followed the Ignore signals differed from the MUA that characterized movement execution (Figure 3; middle-right plots) — similar to the activity that characterized movement inhibition. Moreover, when signal-inhibit trials were compared with ignore-signal trials (Figure 3; lower-right plots), the time at which a significant fraction of sites began to show a difference was delayed by up to 90 ms (mean across sessions ±SE: 48 ±18.3 ms). These findings were confirmed in the other sessions (see Supplementary Fig. 5).

Once we selected sites that participated in inhibitory control — ie, sites that exceeded the thresholding during the SSRT in the previous analysis — we tested whether an inhibitory process developed in the same sites following the Ignore signal by comparing the neuronal activity that was recorded in ignore-signal trials versus no-signal and signal-inhibit trials. The neuronal activity of no-signal trials characterizes the movement generation, whereas that in signal-inhibit trials defines movement suppression. In ignore-signal trials, a movement is generated, and the neuronal activity should be similar to that in no-signal trials. However, if an inhibitory process has been triggered, the ignore-signal activity should bear at least some resemblance to the activity in signal-inhibit trials. The behavioral data and the neuronal population analysis above suggest that this similarity should be observed only when the STD is adopted.

Figure 4 shows that the neuronal patterns expressed after the presentation of either the Stop signal or the Ignore signal are different in the two strategies. In the DTS strategy the only deviation observed is for the activity in the stop signal-inhibit trials. In the STD strategy a more complex pattern emerges: whereas the initial stages of the activities for all 3 types of trials have similar trends, after presentation of the Ignore signal, the activity in ignore-signal trials undergoes modulation that is similar to that in signal-inhibit trials before the end of the SSRT. However, in signal-inhibit trials, subsequently the activity deviated again and displays a pattern similar to that in no-signal trials. The initial strong similarity between the activity in ignore-signal trials and signal-inhibit trials in STD suggests temporary activation of the inhibitory process once a novel signal appears (either Stop or Ignore) after the Go signal.

To provide a representative neuronal dynamics at the population level we adopted a dimensionality reduction approach by principal component analysis (PCA; see Methods). Across strategies, the neuronal activities that were extracted from the 3 trial types had similar evolution until the Stop/Ignore signal presentation, for both monkeys (Figure 5). After these events, the differences between strategies emerged. In both the strategies, the neuronal activity in signal-inhibit trials deviated from the trajectories of no-signal and ignore-signal trials, before the end of the SSRT (the duration is indicated as the thick portion in the trajectories), by traveling toward a subspace in which occupancy was unrelated to movement generation. When monkeys adopted the DTS strategy (Fig. 5 A) there is no evident effect of the appearance of the Ignore-Signal in ignore-signal trials. In contrast, when monkeys adopted the STD strategy (Fig. 5 B), the neuronal dynamics that followed the presentation of the Ignore signal initially underwent a similar evolution as that in signal-inhibit trials, moving toward another region of the space that was unrelated to the movement generation. However, unlike the signal-inhibit trials, after this first deviation, the trajectories shifted again, evolving as in no-signal trials.

**Figure 5:**
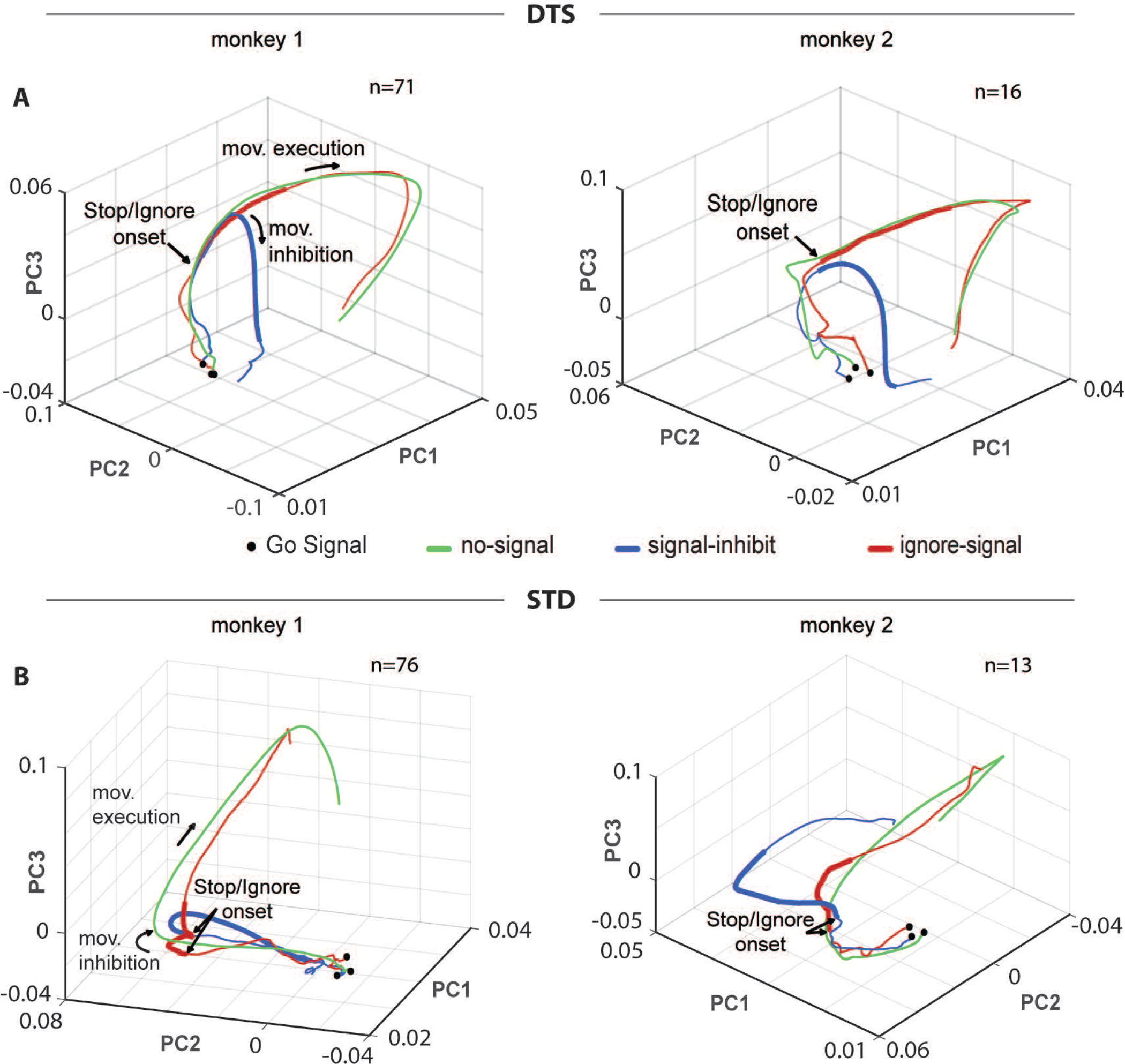
Population neuronal dynamics across animals and behavioral strategies. For each strategy, one session of each monkey is presented. Neuronal trajectories are aligned to the Stop signal for signal-inhibit trials, the Ignore signal for ignore-signal trials, and the hypothetical presentation of the Stop signal in latency-matched no-signal trials, starting from the average presentation of the Go signal (black dots) to 800 ms following the alignment. The thick portions of the trajectories represent the duration of the SSRT.

Once we observed the neuronal dynamics that characterized the various behavioral strategies, we estimated the time that the trajectories deviated after the Stop and Ignore signals. To this end, we evaluated the SNT (Figure 6), defined as the time at which the first derivative of the projection of the neural state onto the single components crossed the zero line (see Methods). The SNT indicates when the trajectories change direction after the Stop or the Ignore signals. We found that the SNT that characterized the inhibition in signal-inhibit trials occurred (mean across sessions ±SE) 70 ±5 ms (DTS strategy) and 82.5 ±10 ms (STD strategy) after the Stop/Ignore signal. There were no signs of deviation that were related to the inhibition in the trajectories in ignore-signal trials when the DTS strategy was adopted.

**Figure 6:**
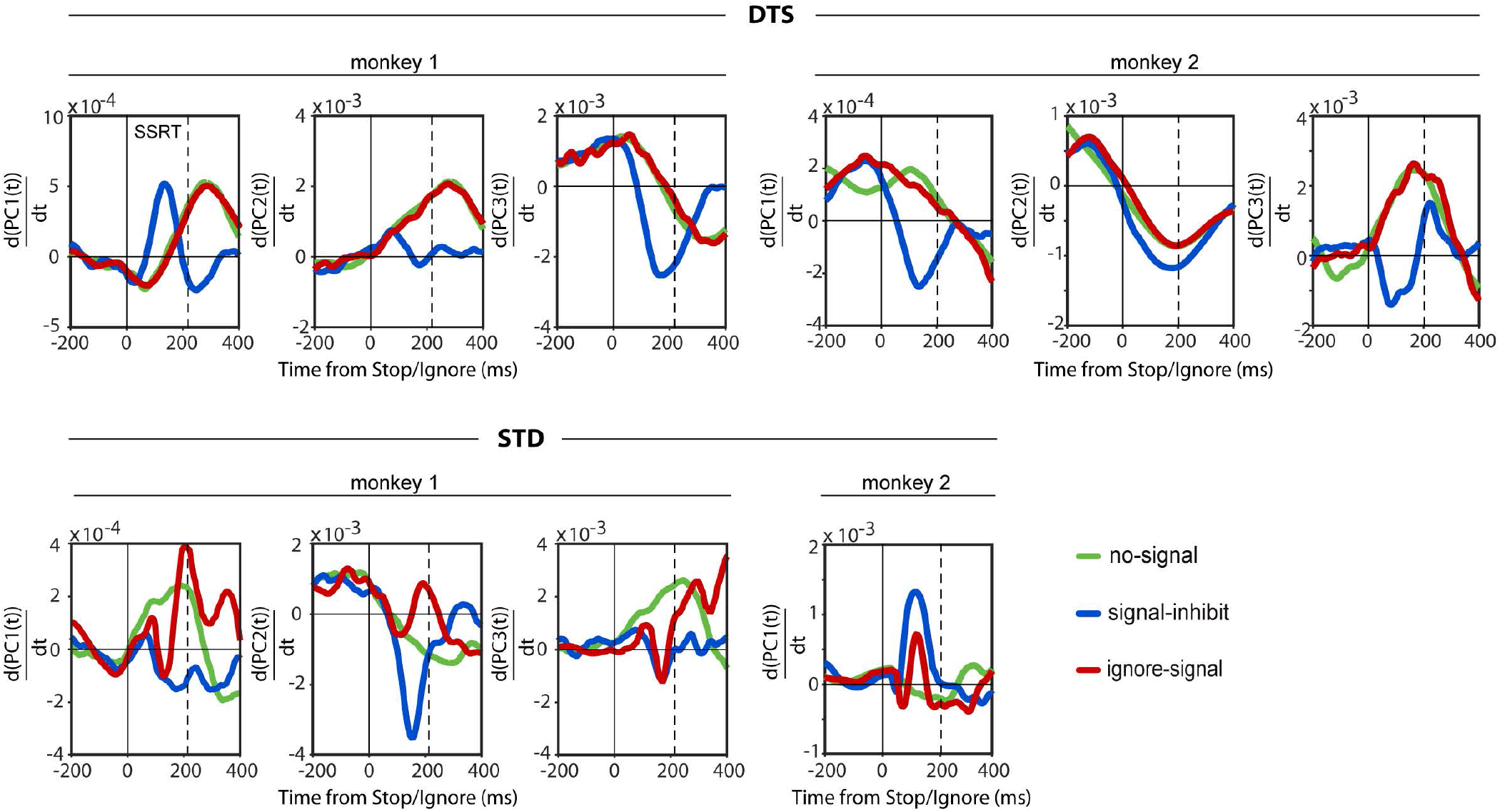
Signal neuronal time across different strategies. First derivatives of the projection of the neuronal state onto the single PCs. Data are from the same sessions in Figure 5

In contrast, when the STD was used (Figure 6, bottom), the derivative in ignore-signal trials crossed the zero value twice after the Ignore signal. The first SNT (mean across sessions ±SE: 96.7 ±14.8 ms) reflects the time at which the trajectories in ignore-signal trials deviated from those in no-signal trials, starting the inversion that characterizes the inhibitory process, whereas the second SNT (mean across sessions ±SE: 158 ±7.8 ms) indicates the end of this temporary inhibition. Based on the difference in time between the 2 SNTs, the duration of this temporary inhibitory process was approximately 60 ms (mean across sessions ±SE: 60.8 ±8.8 ms). See also Supplementary Fig. 6.

Finally, we asked whether it is possible to extract a neuronal signature that allows one to classify the behavioral strategy before the behavioral events that are necessary for its definition occur. We focused on the 200 ms following the Ignore signal. This interval precedes any relevant behavioral event that could be used to describe the strategy. During this window, we measured how the MUA differs when the movement is generated under various conditions (no-signal and ignore-signal trials) with respect to the activity when it is inhibited (signal-inhibit trials). For all of the sessions of the animal with a sufficient number of different sessions for the different strategies (Monkey 1) we performed 2 comparisons: latency-matched no-signal trials vs signal-inhibit trials and ignore-signal trials vs signal-inhibit trials. The trials were selected per the same criteria across sessions (see Methods) and was not influenced by the behavioral strategy. We developed and implemented an algorithm that allowed us to measure whether a difference occurred between the 2 comparisons (see Methods for details). For each comparison, the algorithm extracted a distribution (n=1000) of slopes, every 50 ms at 5-ms increments, of the Euclidean distances between activities — ie, the evolution of the difference between the MUA of 2 trial types over time. The output of the algorithm was a p value — by Kolmogorov-Smirnov test — of the difference in the averaged values of slope distributions (Fig. 7, bottom panels). Two separate sets of p values emerged from the analysis. One set was composed of p values < 0.05 (n=3 sessions, in all cases p<0.0001), and the other set comprised p values > 0.05 (n=2 sessions, in all cases p>0.3). This classification was confirmed by hierarchical cluster analysis (see Supplementary Fig. 7). Then, we traced back to the behavior of the sessions and found that sessions with p value < 0.05 were associated with the STD strategy, whereas those with p > 0.05 correlated with the DTS strategy. Figure 7 summarizes the analysis for 2 representative sessions — 1 for each strategy (for other sessions see Supplementary Fig. 7). For each comparison (latency-matched no-signal trials vs signal-inhibit trials and ignore-signal trials vs signal inhibit trials), the distributions of slopes (top panels) for 2 sample time intervals, t1 (centered at the signal onset) and t2 (centered 150 ms following the signal onset) and the averaged slopes over time (bottom panels) are shown. During the DTS strategy, movement generation was anticipated by a similar evolution of slopes when comparing the MUA in latency-matched no-signal trials and ignore-signal trials with that in signal-inhibit trials, demonstrating that in this case, movement preparation in ignore-signal trials is not influenced by the presence of the Ignore signal.

**Figure 7:**
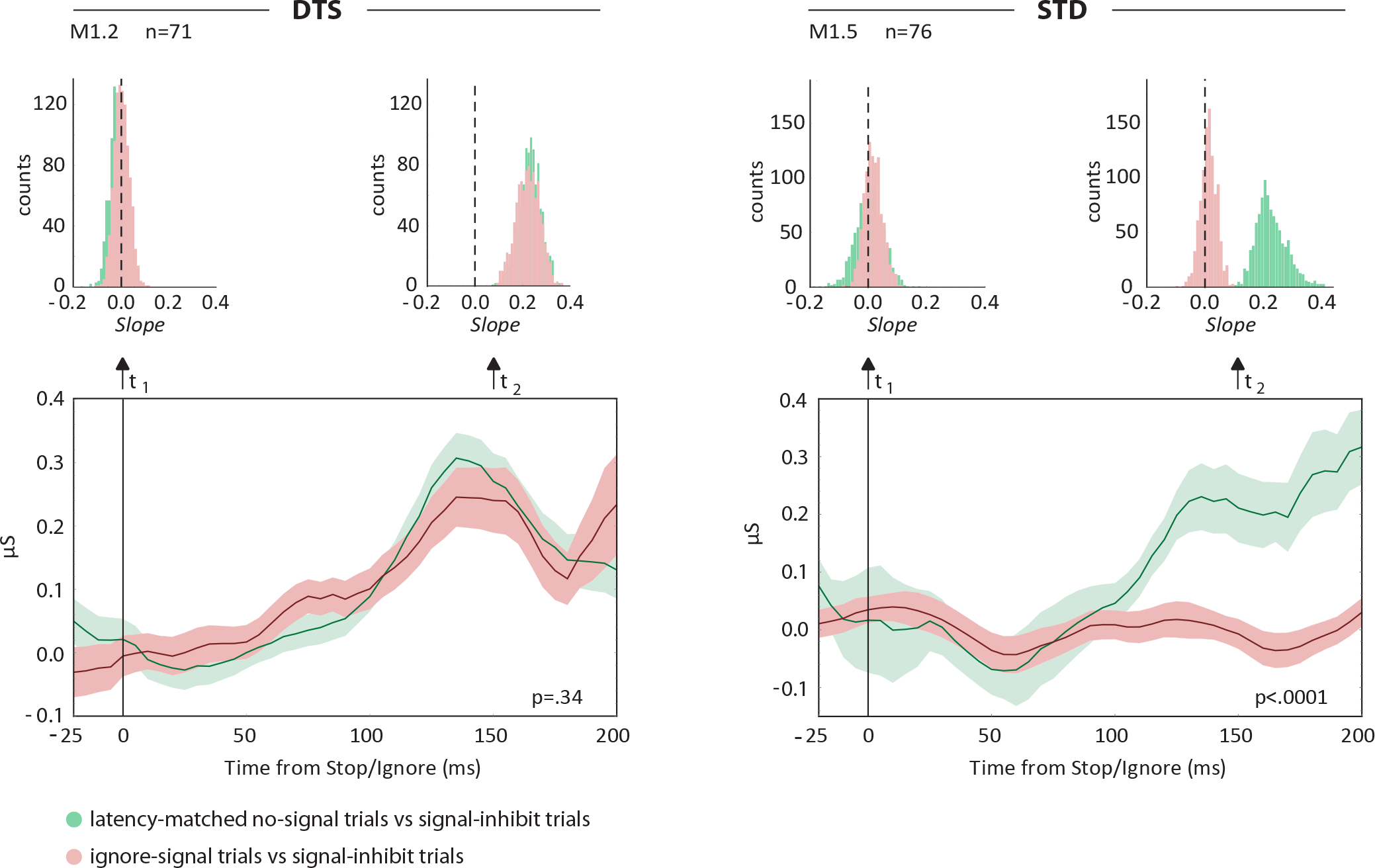
Behavioral strategies are anticipated by specific neuronal modulation following the Ignore signal. Top: For each single strategy, the distributions of slopes of the Euclidian distances of the activity between 2 comparisons are presentedno-signal trials vs signal-inhibit trials and ignore-signal trials vs signal-inhibit trials — for 2 intervals centered to the signal onset (t1) and 150 ms after signal onset (t2). Bottom: Evolution over time of the average slope (μ) of the Euclidean distances for the same comparisons. Data are from the same session in Figure 5, left panels

Conversely, during the STD strategy, movement execution was anticipated by a different evolution of slopes over time. The slopes separated between comparisons at approximately 100 ms after presentation of the signal. After this time, the differences between latency-matched no-signal trials and signal-inhibit trials increase, due to the higher level of motor preparation in latency-matched no-signal trials compared with ignore-signal trials. This greater preparatory activity in latency matched-no signal trials reflects the shorter RTs that will be observed in the DTS.

## 4 Discussion

We examined the neuronal instantiation of inhibition in the PMd using a stimulus selective stop task. This task allowed us to determine the influence of the behavioral strategy on the processing of the Ignore signal. We found that monkeys performed the selective stop task using 2 strategies — STD and DTS — while solving the ignore-signal trials and that there was a strong correlation between the adopted strategy and the effect of the Ignore signal. Specifically, an inhibitory effect on neuronal activity was only observed when the subject implemented the STD strategy. As a result of applying a state-space approach, based on dimensionality reduction, this relationship was congruently established for the single-site- and multisite-level analyses. Consistent with these findings, we assert that the Ignore signal drove the movement inhibition, as represented in the neuronal dynamics of the PMd in a specific behavioral context. In the state-space representation, movement is generated if the neuronal activity travels from a subspace that corresponds to the absence of arm movements to another subspace that will allow their initiation. Along this trajectory, the presentation of a Stop signal can affect the evolution of neuronal activity. In signal-inhibit trials, the neuronal evolution was halted and reversed following the Stop signal: the neuronal activity shifted toward the initial subspace or possibly toward another subspace, the occupancy of which does not allow the initiation of the movement. The emerging behavioral strategies are clearly represented in the neuronal dynamics. When the STD is implemented, in ignore-signal trials, the trajectories are initially affected similarly as in signal-inhibit trials: the neuronal dynamics momentarily reverse their trend, moving toward the initial subspace. However, subsequently, when the meaning of the Ignore signal is clarified, the trajectory follows the evolution that is observed in the no-signal trial, reaching the subspace that leads to initiation of the movement. When DTS strategy is used, the evolution in ignore trials is similar to that for no-signal trials: in this case, there is no evidence of the activation of the inhibitory process. The observation that the initiation of arm movement requires a shift from one subspace to another is consistent with findings that were obtained with the reaching delay task and a dynamic system approach (Shenoy et al., 2013; Kaufman et al., 2014; Ames et al., 2014). Several of these studies showed that movements are prepared during the delay epoch, in a subspace (output-null dimension) that prevents neuronal activity from affecting the muscle and generating movements. Movements are executed when neuronal trajectories reach another subspace (output-potent dimension), in which rotational dynamics appear (Elsayed and Cunningham, 2017; Churchland et al., 2012). In our study, the neuronal activity remained in a subspace during the first period of all the trials and during the last period of the signal-inhibit trials, when movements are not generated. As shown by the derivative analysis, when the STD is adopted in ignore-signal trials, the neuronal activity travels first toward the movement initiation subspace, reverses slightly, and then finally re-reverses to allow the movement generation. This finding is the first demonstration of the dependence of movement inhibition on the adopted strategy. Following presentation of the Ignore signal, clear dynamics of inhibition emerged only in the STD strategy, correlating directly with behavior and occurring before the behavioral strategy could be established.

Although we examined the neuronal dynamics that underlie movement inhibition using a state-space approach, the study of the neural basis of movement inhibition for other systems and structures has been oriented primarily toward the characterization of specific cell types. The combination of countermanding task and neurophysiology has yielded significant results for the saccadic system in monkeys and for the function of basal ganglia in movement inhibition in rats. In the monkey saccadic system, movement control has been ascribed to 2 cell types in the frontal eye fields (FEF) and superior colliculus (SC): movement cells and fixation cells. Saccades are made when movement cells increase their firing activity, whereas fixation cells decrease it; saccades are inhibited when the opposite pattern occurs, following the Stop signal and before the end of the SSRT (Hanes et al., 1998; Paré and Hanes, 2003). A recent study found that a subpopulation of dopaminergic neurons in the substantia nigra (SNc) and a connected subpopulation of striatal neurons fire differently when saccades are made versus withheld and before the end of the SSRT (Ogasawara et al., 2018). However, the relationship between nigro-striatal, cortical, and collicular activities must be clarified. Clear neuronal types that participate in various aspects of movement inhibition have also been found in the basal ganglia of rats: a series of studies by Berke and colleagues (Schimdt et al., 2013; Mallet et al., 2016) have found that movement inhibition can be mapped to different cell types and structures. When a Stop signal occurs, movements are first paused briefly by the neuronal activity in the subthalamic nucleus and substantia nigra and then cancelled, if necessary, by neurons in the pallidus, affecting the striatum (Schmidt and Berke, 2017). This modulation purportedly affects the neuronal dynamics in cortical motor regions. It is not possible to establish the definitive relationship between neuronal types and aspects of movement inhibition for the limb cortical motor system in primates (Kaufman et al., 2010). Single cells in the PMd show are heterogeneous, preventing any simple or mechanistic classification, as can be performed for the saccadic system and basal ganglia (Kaufman et al., 2010; Mirabella et al., 2011). In our study, we could not execute this classification, because we recorded the spectral derived MUA, which the reflects spiking activity of small population of neurons that surround the tips of the electrodesby their very nature, many neuronal types can contribute to the activity. However, the patterns that we observed strongly resemble the typical patterns of activity in single cells, and most importantly, their function in movement inhibition is supported by their SNTs before the SSRT, as observed in other studies (Pani et al., 2018). The system-level implementation of movement inhibition requires communication between various regions, each of which can experience specific neuronal implementation of the inhibition process, by specific neural type or population code (Aron et al., 2010; Pouget et al., 2017). Assuming that primates use a basal ganglia-based mechanism for control of limb movements, similar to that in rodents, the modulations during movement inhibition that are recorded in the basal ganglia (and in other regions) might appear to be heterogeneous when viewed at the motor cortical level (Oldenburg and Sabatini, 2015; Mattia et al., 2013), thus rendering the state-space approach a suitable method for describing the inhibition of limb movements.

This study strengthens the evidence in favor of the PMd as a site of movement control. It is well established that the PMd continuously signals the momentary decision state about forthcoming movements (Thura and Cisek, 2014; Kaufman et al., 2014) and movement parameters (velocity, reaction time). The data that support its function in movement inhibition, as required by the stop-signal task, are accumulating (Mirabella et al., 2011; Pani et al., 2013; Pani et al., 2018). In this study, we demonstrated that the function of the PMd in inhibition is strategy-dependent — ie, the PMd reflects movement inhibition only when it is behaviorally relevant. The presentation of the Ignore signal does not drive the inhibition *per se* but only under the conditions in which the signal can influence the movement plan. Further, by using the stop task, movement inhibition-related activity is clearly represented in PMd neurons (Mirabella et al., 2011; Pani et al., 2018), whereas attempts to detect coherent activation in other areas that control limb movements have been unfruitful (Scangos and Stuphorn, 2010).

In the literature on selective inhibition in humans (Bissett and Logan, 2014), the DTS strategy is usually described as Independent DTS to distinguish it from a Dependent DTS strategy. The Dependent DTS strategy is characterized by signal-respond RTs that are no slower than no-signal RTs, thus violating the race-model independence assumption. In this last case, some form of inhibition is believed to occur through the trial, delaying the response in respond stop trials (Bissett and Logan, 2014; Sebastian et al., 2017). We did not find any behavioral evidence that could be referred to as Dependent DTS. One possible reason is that the monkeys were highly trained in performing the task and that the amount of training led them to develop a more efficient strategy in deciding between stopping and moving. However, the STD and DTS are also observed in humans (Sebastian et al., 2017; Bissett and Logan, 2014). One additional caveat is that in the STD, the SSRT should be shorter than when the DTS is used (Bissett and Logan, 2014). This difference is related to the presence of a longer discrimination stage following the Ignore signal in the Independent DTS. We did not observe such an effect after analyzing our data: in general, the length of the SSRT did not differ between strategies. Further, analogous results have been observed in humans by similar studies (Bissett and Logan, 2014; Sebastian et al., 2017). Thus, qualitatively and strategically, performance of monkeys and humans is alike, and our data can be used to hypothesize similar neuronal dynamics in humans.

