## Supplemental files for "Neuronal dynamics of signal selective motor plan cancellation in the macaque dorsal premotor cortex"

[§]=corresponding author

### Supplementary materials

**Supplementary Figures: 7**

**Supplementary Tables:1**

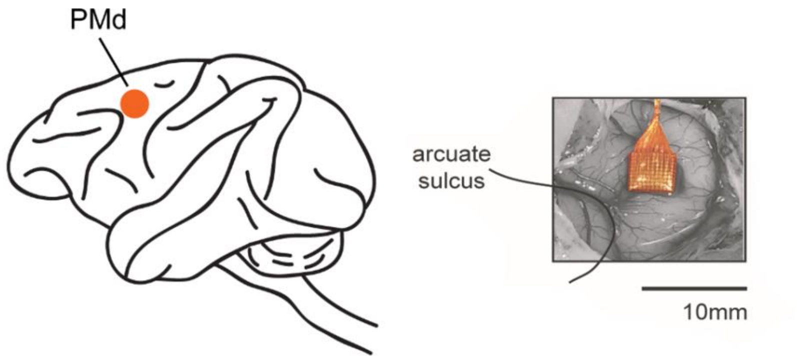


**Supplementary Figure 1**. Site of recordings. Left: approximate location for both animals. Right: example of picture taken during surgery; Monkey 1.

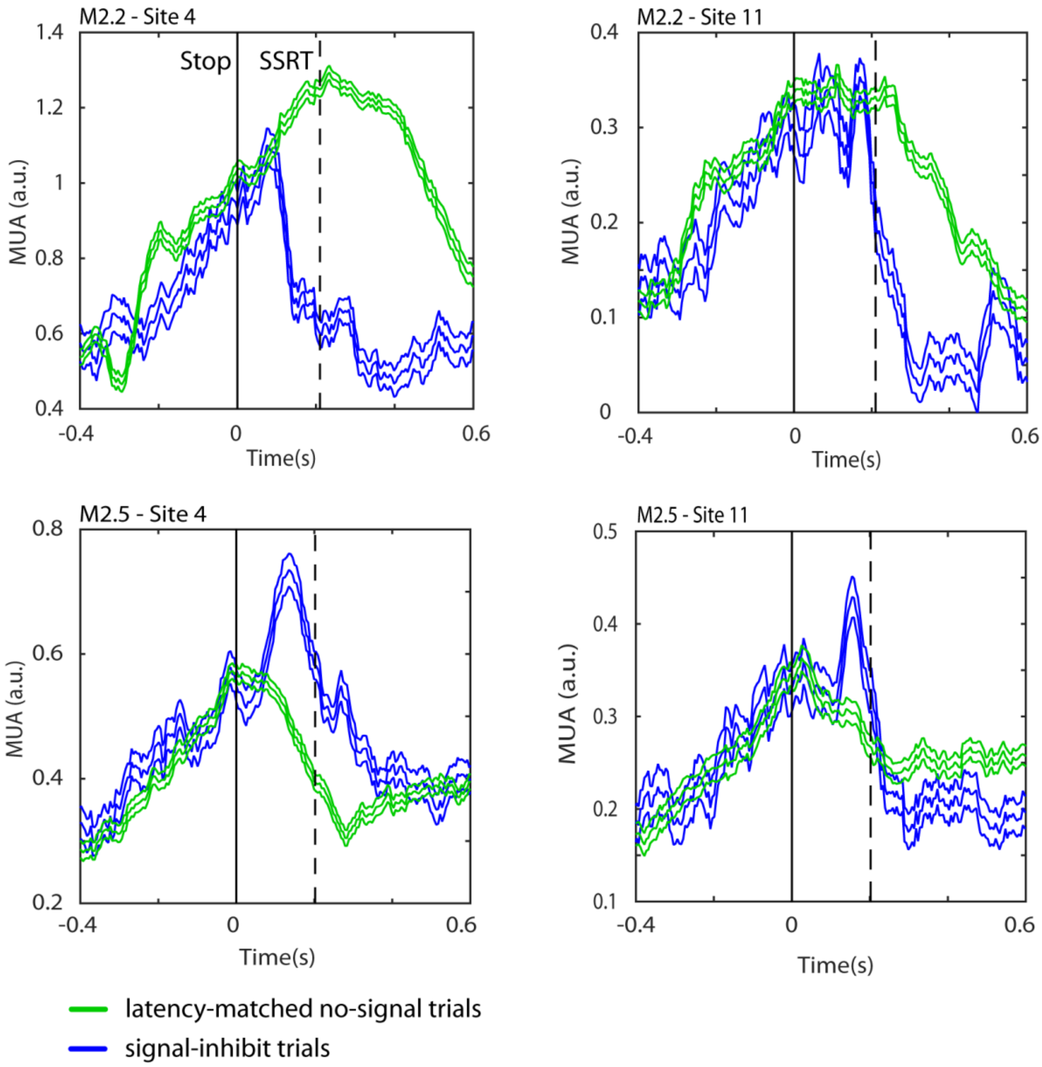


**Supplementary Figure 2.** Different patterns of average MUA(±SE) recorded from the same sites in two separate sessions (M2.2, M2.5). These are the selected sites shared by the two sessions that we merged for the PCA (See Fig. 5; DTS strategy-monkey 2).

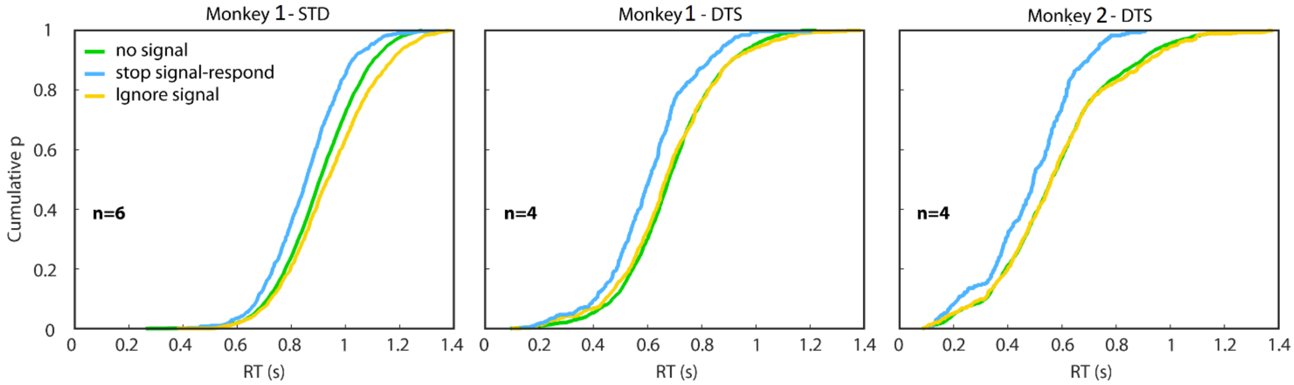


**Supplementary Figure 3**. RTs in no-signal, stop-signal-respond and ignore-signal trials with data collapsed across sessions for different behavioral strategies. For data from the single STD session in monkey 1 see Supplementary Table 1.

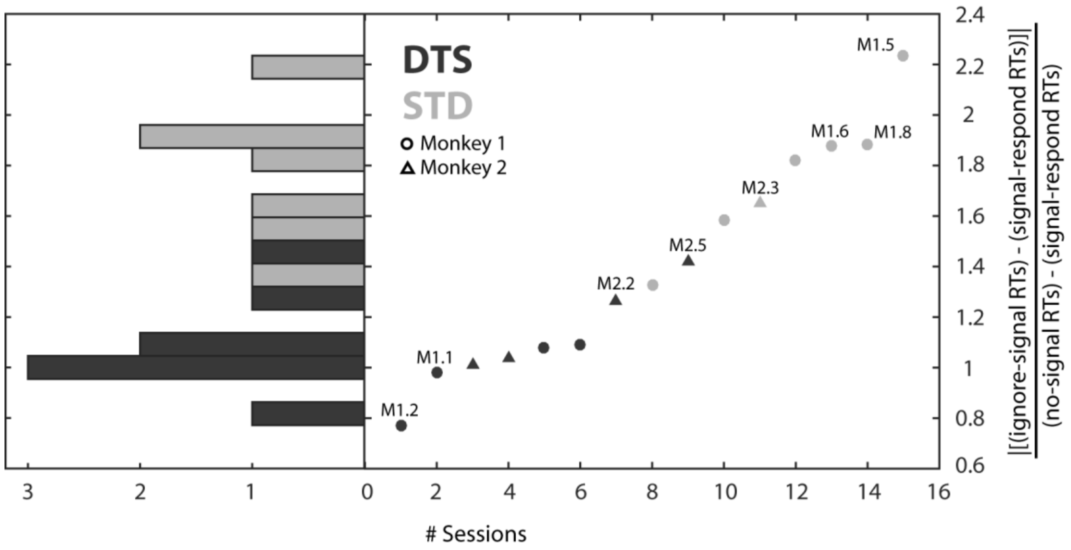


**Supplementary Figure 4**. Distribution of metrics representing the strategies. We calculated, for each behavioral session, the ratio between the absolute value of the signal-respond RTs minus the ignore-signal RTs, and the no-signal RTs minus signal-respond RTs. The left panel shows the histograms of the metrics obtained (DTS, black bars; STD, grey bars), while the right panel shows the ranked values of the same metrics. Colours represents the strategy assigned on the basis of the typical methods adopted in literature (see main text). Sessions employed for neuronal analysis are made explicit above the symbols (e.g., M1.1. session 1 for Monkey 1 as in table 1).

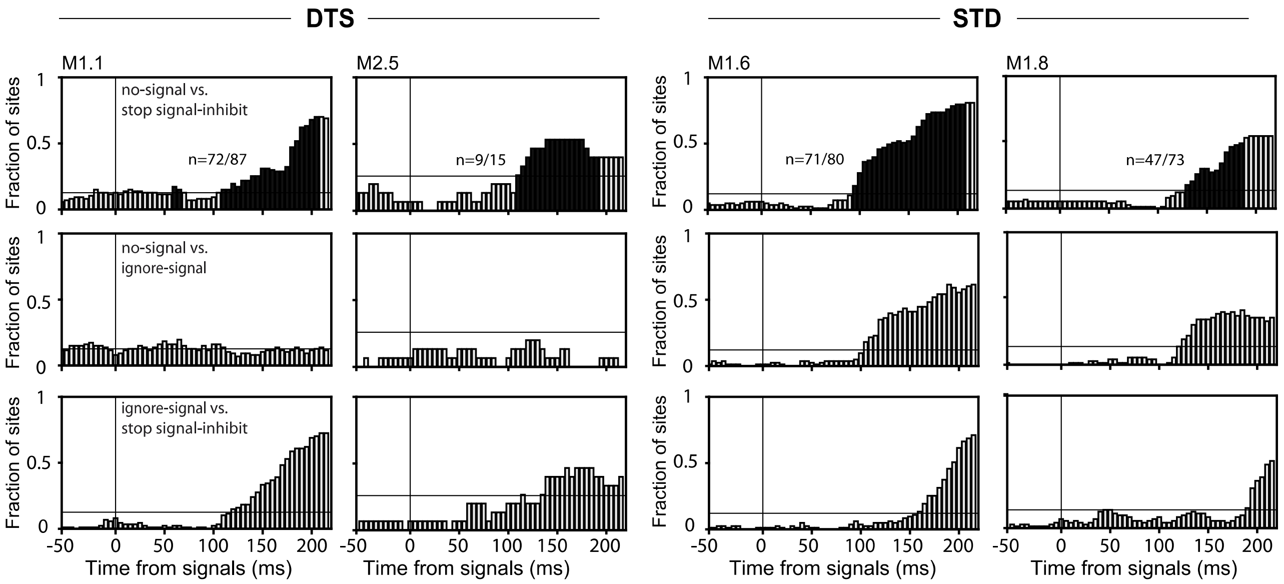


**Supplementary Figure 5**. Neuronal characterization of behavioral strategies at the population level, further example sessions. Fraction of sites whose MUA significantly differed between the trial types under comparison, separately for each monkey and under the two strategies (DTS and STD). Other conventions and symbols as in Figure 3 of the main text. Sessions employed are made explicit above each top panel.

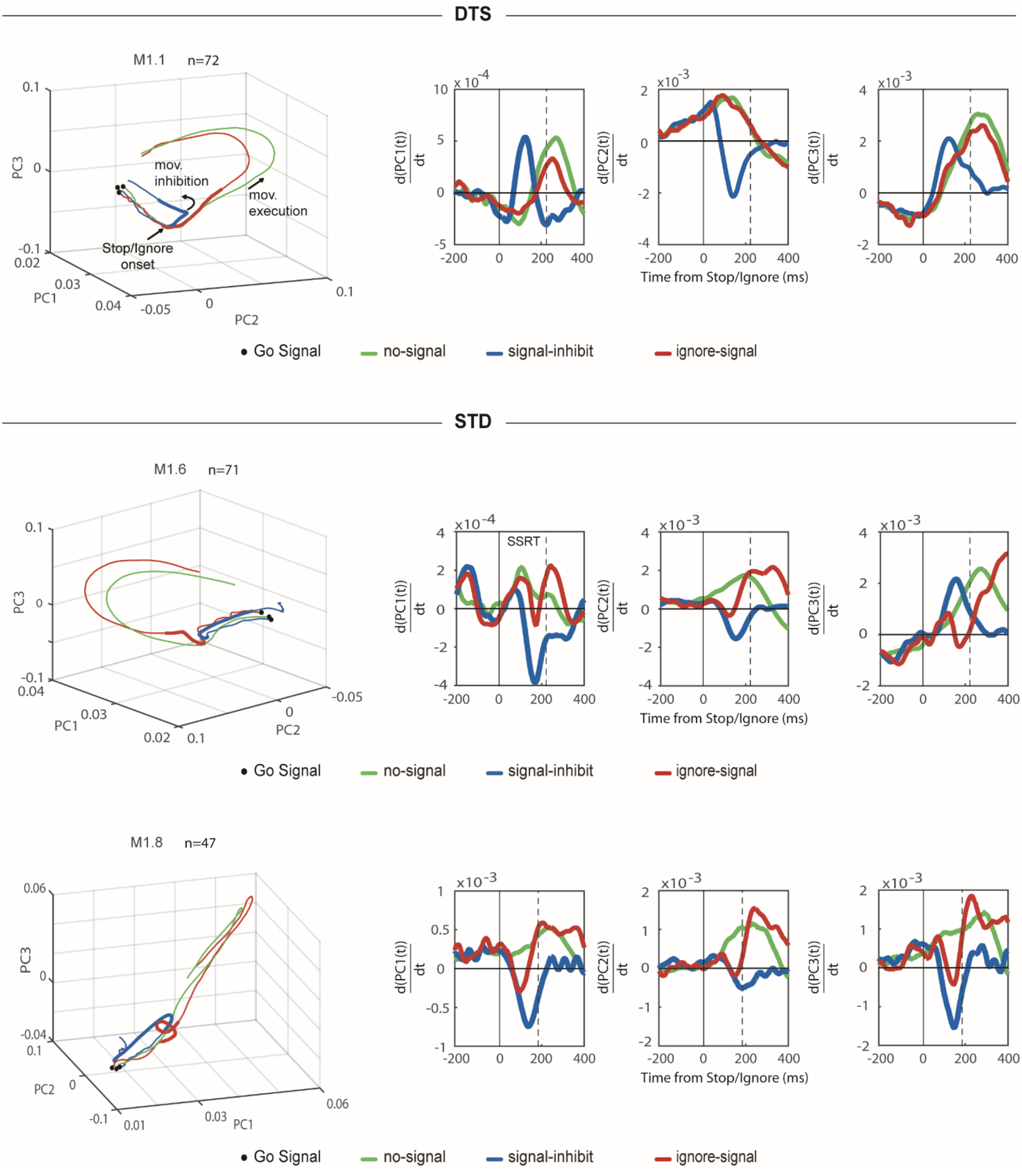


**Supplementary figure 6** Neural dynamics and SNT computation in the different trial types for three additional sessions of monkey 1 (one for the DTS strategy and two for the STD strategy). Other conventions and symbols as in Figure 6 of the main text.

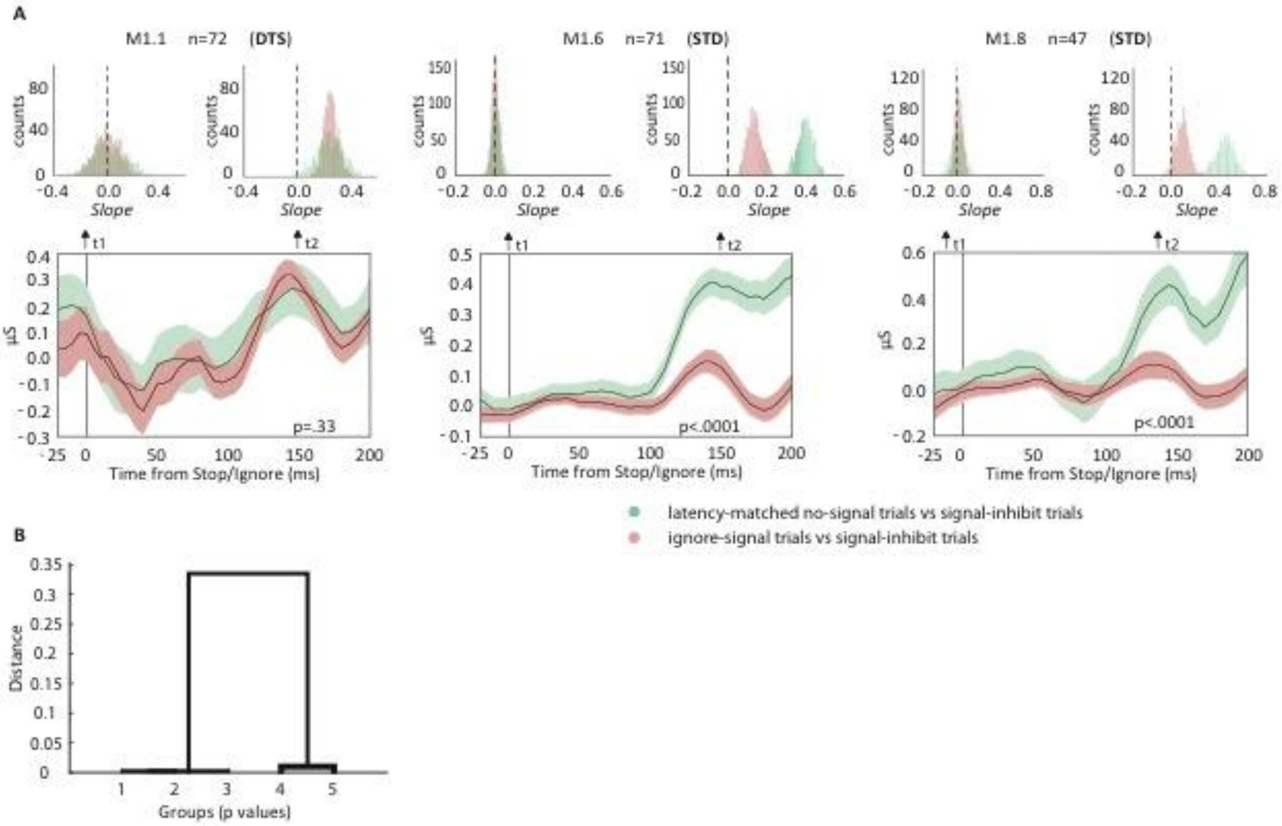


**Supplementary Figure 7**. **A**) Analysis performed to classify the behavioral strategies based on the neuronal activity for three additional sessions. Conventions and symbols as in Figure 7 of the main text **B**) Hierarchical cluster tree based on the Euclidian distances between the observed p values and a threshold of p=0.05 (cophenetic correlation coefficient = 0.99). One cluster was composed by three sessions associated with p value<0.05 (in all cases p<0.0001) from the STD strategy. The second cluster was composed by two sessions with p value>0.05 (in all cases p>0.3) from the DTS strategy.

| Monkeys Sessions | n trials | Fore-period delay range (50ms step) | no-signal RT Mean(SD) | signal-respond RT Mean(SD); p | ignore-signal RT Mean(SD); p | P(Resp) | median SSD (s) | SSRT (ms) | Strategy |
|---|---|---|---|---|---|---|---|---|---|
| M1.1* | 751 | 800-1150 | 649(144.4) | 542.3(128.8); p<.001 | 646.7(83.2); p=.87 | 0.45 | 0.420 | 218 | **DTS** |
| M1.2* | 1016 | 800-1150 | 595.4(187.5) | 492(155); p<.001 | 571.8(216.5); p=.13 | 0.39 | 0.320 | 223 | **DTS** |
| M1.3 | 756 | 800-1150 | 657.9(145.8) | 555.8(125); p<.001 | 667.3(152.4); p=.5 | 0.41 | 0.420 | 192 | **DTS** |
| M1.4 | 894 | 0 | 844.7(120.2) | 808.7(120.6); p=.01 | 874.3(194.3); p=.004 | 0.46 | 0.620 | 215 | **STD** |
| M1.5* | 1210 | 0 | 819.5(117) | 785.8(135.2); p=.005 | 861.1(168.3); p<.001 | 0.47 | 0.620 | 230 | **STD** |
| M1.6* | 1311 | 0 | 908.6(132.7) | 853.8(136.9); p<.001 | 956.8(141.4); p<.001 | 0.48 | 0.720 | 213 | **STD** |
| M1.7 | 1457 | 320-570 | 957(160.3) | 871.3(157); p<.001 | 985.2(142); p=.01 | 0.47 | 0.721 | 213 | **STD** |
| M1.8* | 1336 | 320-570 | 968.6(159.6) | 935.4(149.1); p=.02 | 998(146.5); p=.01 | 0.48 | 0.720 | 194 | **STD** |
| M1.9 | 1375 | 320-570 | 908(143.8) | 841.8(108.5); p<.001 | 946.6(144.7); p<.001 | 0.46 | 0.720 | 223 | **STD** |
| M1.10 | 1311 | 0 | 786.1(154) | 727.4(132.2); p<.001 | 791.1(176.2); p=.67 | 0.46 | 0.620 | 188 | **DTS** |
| M2.1 | 752 | 800-1150 | 659.4(209.4) | 558(144); p<.001 | 664(223.8); p=.6 | 0.48 | 0.420 | 225 | **DTS** |
| M2.2* | 828 | 800-1150 | 679.6(192.4) | 590.5(130.2); p<.001 | 703.7(214.5); p=.2 | 0.46 | 0.420 | 209 | **DTS** |
| M2.3* | 1045 | 800-1150 | 600.2(149) | 544.2(90.6); p<.001 | 637(174.7); p=.003 | 0.49 | 0.420 | 203 | **STD** |
| M2.4 | 497 | 800-1150 | 583(189.5) | 508(117.2); p=.016 | 615(191.1); P=.16 | 0.44 | 0.320 | 182 | **DTS** |
| M2.5* | 944 | 800-1150 | 405(149) | 337(121.3); p<.001 | 406(169); P=.96 | 0.56 | 0.320 | 202 | **DTS** |

**Table 1**. Details of behavioral performance from all sessions. (*) Sessions included in the neuronal analysis.
